# Speech Fine Structure Contains Critical Temporal Cues to Support Speech Segmentation

**DOI:** 10.1101/508358

**Authors:** Xiangbin Teng, Gregory B. Cogan, David Poeppel

## Abstract

Segmenting the continuous speech stream into units for further perceptual and linguistic analyses is fundamental to speech recognition. The speech amplitude envelope (SE) has long been considered a fundamental temporal cue for segmenting speech. Does the temporal fine structure (TFS), a significant part of speech signals often considered to contain primarily spectral information, contribute to speech segmentation? Using magnetoencephalography, we show that the TFS entrains cortical oscillatory responses between 3-6 Hz and demonstrate, using mutual information analysis, that (i) the temporal information in the TFS can be reconstructed from a measure of frame-to-frame spectral change and correlates with the SE and (ii) that spectral resolution is key to the extraction of such temporal information. Furthermore, we show behavioural evidence that, when the SE is temporally distorted, the TFS provides cues for speech segmentation and aids speech recognition significantly. Our findings show that it is insufficient to investigate solely the SE to understand temporal speech segmentation, as the SE and the TFS derived from a band-filtering method convey comparable, if not inseparable, temporal information. We argue for a more synthetic view of speech segmentation – the auditory system groups speech signals coherently in both temporal and spectral domains.

## Introduction

Parsing the continuous speech stream into appropriate units for subsequent perceptual and linguistic analyses serves as the basis for recognition (Poeppel 2003; Ghitza 2012; Giraud and Poeppel 2012). Speech is typically argued to be comprised of slower amplitude fluctuations, called the speech amplitude envelope (SE) and faster temporal and frequency events, called the temporal fine structure (TFS). These distinct aspects of speech signals have been investigated in different experiments to elucidate their relative contributions (Smith et al. 2002; Xu and Pfingst 2003; Zeng et al. 2004; Lorenzi et al. 2006; Moore 2008; Shamma and Lorenzi 2013; Ewert et al. 2018). Evidence suggests that the human auditory system relies on temporal information supplied by the SE for grouping speech information (Ghitza and Greenberg 2009; Ghitza 2012; Ding and Simon 2014). Behavioural data demonstrate that four bands of noise modulated by the SE suffice for speech recognition (Shannon et al. 1995). Neurophysiological evidence shows that cortical entrainment to the SE shows a high correlation with speech intelligibility (Luo and Poeppel 2007; Ding and Simon 2012; Giraud and Poeppel 2012; Peelle et al. 2013; Doelling et al. 2014). Speech, though can be decomposed into two parts (SE and TFS) using a ‘filterbank’ method (Shannon et al. 1995; Smith et al. 2002), containing coherent amplitude fluctuations and spectral cues. Does the auditory system primarily exploit the SE for segmentation or use additional attributes of the speech signal when segmenting the incoming speech stream?

The TFS is argued to convey distinct cues not present in the SE (Gilbert et al. 2007) and to play an important role in mediating speech perception in challenging backgrounds (Hopkins et al. 2008; Moore 2008; Swaminathan et al. 2016). It was demonstrated in neurophysiological studies that the TFS contributes to robust cortical entrainment for speech in noise (Ding and Simon 2014). Indeed, after eliminating amplitude fluctuations, the processed speech signal can still entrain cortical oscillatory responses (Zoefel and VanRullen 2015; 2016). These studies argue for an important role played by the TFS, which complements the traditional view that the SE is the dominant cue for speech segmentation. However, it remains unclear how the TFS contributes to robust cortical entrainment to speech and whether the TFS and SE play different roles in speech segmentation.

Various studies suggest that the envelope can be ‘recovered’ from the TFS through cochlear processing (Ghitza 2001; Zeng et al. 2004; Shamma and Lorenzi 2013). From a neurophysiological perspective, it is argued that cues in the spectral structure of speech (without amplitude modulations) entrain cortical oscillations and provide temporal information (Zoefel and VanRullen 2015). The TFS provides acoustic cues to help form auditory objects for grouping and entrainment (Ding and Simon 2012; Ding et al. 2014). But the cues, whether acoustic or high-level, are not clearly specified in previous studies, and it is not well understood how these cues are extracted from the TFS.

Here we first take a traditional approach by separating the SE and the TFS using the ‘filterbank’ method and test whether the human auditory system can capitalize on the TFS to temporally segment speech signals. If the TFS and the SE represent different aspects of speech signals and the auditory system primarily extracts temporal information from the SE for segmenting speech, as indicated by the previous studies (Luo and Poeppel 2007; Ghitza and Greenberg 2009; Ding and Simon 2012; Ghitza 2012; Giraud and Poeppel 2012; Peelle et al. 2013; Ding and Simon 2014; Doelling et al. 2014), we would expect that the TFS alone cannot entrain cortical oscillatory responses (to the same level) as the SE does. In contrast, if we do find that the TFS supplies sufficient temporal information and robustly entrains cortical oscillatory responses, we aim to determine how temporal information is extracted from the TFS, as well as the nature of this temporal information. We then test behaviourally whether the TFS will provide temporal cues and help increase intelligibility when the SE is disrupted temporally.

We show through neurophysiological results that the TFS, similar to the SE, robustly entrains cortical oscillatory responses. The patterns of cortical entrainment differentiate between the TFS of different sentences. To better understand what in the TFS elicits robust cortical entrainment, we modified a method - cochlear scaled correlation (Stilp and Kluender 2010) - to derive temporal information from the TFS. We compute the mutual information between neurophysiological responses evoked by the TFS and 1) the original SE, 2) the recovered envelope from the TFS, and 3) the derived temporal information through spectral correlation in the TFS. We determine that the temporal information in the TFS is highly relevant to the SE, and is contributed by the spectral correlation of the TFS as well as by the recovered envelope. We further show that spectral resolution strongly affects the ability to extract temporal information from the TFS. Next, we temporally distort speech by using a widely cited but not often-used manipulation, locally reversing speech segments (Saberi and Perrott 1999; Kiss et al. 2008; Stilp et al. 2010). We demonstrate that the TFS helps restore critical temporal information and significantly improves intelligibility of temporally distorted speech.

Our results demonstrate that the TFS provides significant temporal information for segmenting speech. The TFS and the SE convey comparable, if not inseparable, temporal information for speech segmentation. We conclude that speech segmentation and cortical entrainment to speech are a result of tracking both the temporal and spectral structure of speech.

## Materials and Methods

### Ethics statement

The study was approved by the New York University Institutional Review Board (IRB# 10–7277) and conducted in conformity with the 45 Code of Federal Regulations (CFR) part 46 and the principles of the Belmont Report. All the participants in Experiment 1 and 2 gave informed written consent.

### Participants

Fifteen Chinese native speakers from New York University took part in Experiment 1. The data from three participants were excluded from MEG analysis because trial orders were not recorded during neurophysiological recording. One participant was further excluded from the mutual information analysis because part of data from this subject was removed after preprocessing due to a noise issue, which resulted in the loss of the trial order for the mutual information analysis. Therefore, in Experiment 1, the analysis included the neurophysiological data from 12 participants for inter-trial phase coherence and single-trial classification (6 females; age ranging from 21 to 32; right-handed) and from 11 participants for the mutual information analysis (6 females; age ranging from 21 to 32; right-handed). Handedness was determined using the Edinburgh Handedness Inventory (Oldfield 1971).

Twenty-One Chinese native speakers studying at New York University took part in Experiment 2: ten in Experiment 2A (5 females; age ranging from 23 to 35; all self-reported right-handed) and eleven in Experiment 2B (8 females; age ranging from 22 to 28; all self-reported right-handed). One participant was excluded from Experiment 2B because of poor performance. All participants had normal hearing and no neurological deficits according to their self-report.

### Stimuli

One hundred Chinese sentences from the Mandarin Hearing in Noise Test were used in the present study (Fu et al. 2011; Zhu et al. 2014). All sentences are composed of 7 syllables each and have similar grammatical structure and are spoken by a female speaker. The overall power of all sound files was normalized to the same value before further acoustic processing. We first decomposed the sentences into envelopes and TFS (see detailed method also in (Smith et al. 2002)). We filtered the speech signal into 16 bands using cochlear filter banks spanning from 80 to 8820 Hz and created analytic signals for each frequency band through a Hilbert transformation (see Figure 1A for an illustration). The envelope is computed as the magnitude of the analytic signal and TFS was reconstructed by applying a cosine function to the phase series of the analytic signal for each frequency band. We selectively chose 16 bands for decomposing the speech signal because previous studies show that the envelope cannot be recovered from the TFS, or the recovered envelope from TFS is no longer beneficial for speech recognition when 16 bands are used (Gilbert and Lorenzi 2006; Sheft et al. 2008), which enables us to further investigate what else in the TFS, besides the recovered envelope, can contribute to speech segmentation.

**Figure 1.**
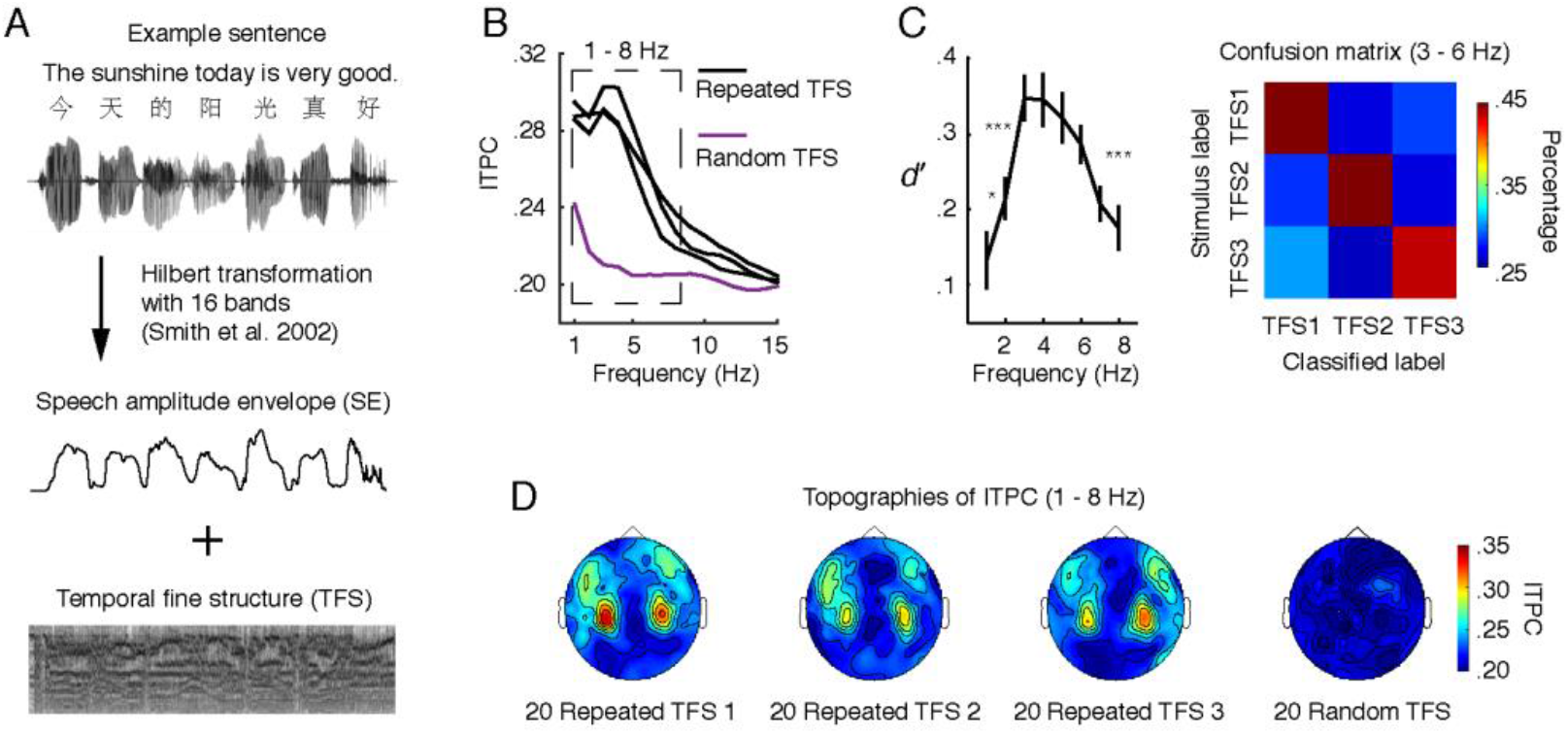
Cortical entrainment to TFS. (A) Speech signals were decomposed using 16 bands into the speech amplitude envelope (SE) and the temporal fine structure (TFS). (B) The repeated TFS stimuli evoke robust cortical entrainment from 1 to 8 Hz. X-axis: frequency of neural response. Y-axis: inter-trial phase coherence (ITPC). Black lines show entrainment to the three repeated TFS stimuli, separately. The violet line shows cortical entrainment to 20 random TFS sentences, used as a baseline for the ITPC. The dashed box indicates frequencies where ITPC of the repeated TFS stimuli is significantly larger than ITPC of the 20 random TFS stimuli (*p* < 0.05, paired t test, Bonferroni corrected). (C) Left panel, classification analysis conducted on the phase series within each frequency band. The results show that the neural signals in frequencies between 3 and 6 Hz are the most informative to separate out different repeated TFS stimuli. Right panel: the group-averaged confusion matrix between 3 and 6 Hz. Each repeated TFS stimulus can be robustly classified. (D) Topographies for ITPC between 1 and 8 Hz confirm that the cortical entrainment to repeated TFS stimuli is of auditory origin. The error bars represent +/- SEM over subjects. Asterisks show significant level (***, *p* < 0.001; *, *p* < 0.05).

In Experiment 1, we generated 28 TFS stimuli reconstructed from twenty-eight randomly selected sentences by averaging TFS of each frequency band across 16 bands for each sentence.

In Experiment 2, we processed the sentences and created four types of reversed speech: directly reversed speech (R), envelope reversed speech (ER), fine structure reversed speech (FSR), and envelope reversed noise-vocoded speech (ERNV). To generate R sentences, we cut each sentence into short segments using a rectangular window (e.g. 50 ms) and reversed each segment temporally. Then we concatenated the reversed segments of each sentence in the original order to form a new sentence, whose local segments were temporally reversed. The R sentences were generated in the same way as in previous studies (Saberi and Perrott 1999; Stilp et al. 2010).

Figure 3A illustrates procedures to generate ER, FSR, and ERNV sentences. We cut the envelope of each band into segments using a fixed window size and reversed each segment temporally, and then concatenated them to form a reversed envelope for each band. The new reversed envelope was then used to modulate the intact TFS of the corresponding band to generate the ER sentences. We used the new reversed envelopes to modulate narrow band noise of corresponding frequency bands to get the ERNV sentences. We kept the envelopes intact while cutting TFS into segments and reversing TFS segments to form reversed TFS. The intact envelopes and the reversed TFS were put together to form FSR sentences.

In Experiment 2A, R, ER, and FSR sentences were used to test intelligibility on each type of speech. Six window sizes were used to cut speech into segments for the R sentences: 30, 50, 70, 80, 90, and 120 ms; six window sizes for the ER sentences: 30, 70, 90, 120, 150, and 200 ms; three window sizes for the FSR sentences: 30, 150, and 300 ms. In Experiment 2B, ER and ERNV sentences were used. Six window sizes were chosen for the ERNV sentences: 30, 50, 70, 80, 90, and 120 ms; six window sizes for the ER sentences: 30, 70, 90, 120, 150, and 200 ms.

All stimuli used in Experiment 1 and 2 were normalized to ~65 dB SPL. The stimuli were delivered through plastic air tubes connected to foam ear pieces (E-A-R Tone Gold 3A Insert earphones, Aearo Technologies Auditory Systems) in Experiment 1 and through Sennheisser 370 headphones in Experiment 2.

### Experiment 1: MEG procedure

We selected TFS stimuli of 28 total sentences. 25 TFS stimuli of different sentences were presented once each and 3 TFS stimuli of 3 different sentences were presented 25 times each. All TFS stimuli were pseudo-randomly presented in one block during MEG recordings. To keep subjects alert, after hearing each TFS stimulus participants were prompted to make a judgment via a button box on whether TFS stimuli sounded like speech or not. The behavioral responses were not analyzed because all participants reported that TFS stimuli did not sound like speech. This could be because the TFS was derived using 16 bands and the envelope cues cannot be recovered from the TFS (Smith et al. 2002; Hopkins et al. 2010). The participants had no prior knowledge on the TFS stimuli or on the sentences from which the TFS stimuli were derived. The inter-trial interval (ISI) of 1.5 – 2 s began after the key press. The ISI was used as a baseline for MEG analysis.

### MEG recording and preprocessing

MEG signals were measured with participants in a supine position, in a magnetically shielded room using a 157-channel whole-head axial gradiometer system (KIT, Kanazawa Institute of Technology, Japan). A sampling rate of 1000 Hz was used with an online 1-200 Hz analog band-pass filter and a notch filter centered around 60 Hz. After the main experiment, participants were presented with 1 kHz tone beeps of 50 ms duration as a localizer to determine their M100 evoked responses, which is a canonical auditory response (Roberts et al. 2000). 20 channels with the largest M100 response in both hemispheres (10 channels in each hemisphere) were selected as auditory channels for each participant individually. Further analysis was conducted only on the selected channels.

MEG data analysis was conducted in MATLAB using the Fieldtrip toolbox (Oostenveld et al. 2011) and the wavelet toolbox. Raw MEG data were noise-reduced offline using the time-shifted PCA (de Cheveigné and Simon 2007) and sensor noise suppression (de Cheveigné and Simon 2008) methods. A low-pass filter with cutoff frequency of 100 Hz was applied offline on the de-noised data in the MEG160 software (Yokogawa Electric Corporation and Eagle Technology Corporation, Tokyo, Japan) and the preprocessed MEG data was then downsampled to 500 Hz. Trials were visually inspected, and those with artifacts such as channel jumps and large fluctuations were discarded. An independent component analysis was used to correct for eye blink-, eye movement-, heartbeat-related and system-related artifacts. Each trial was divided into 6s epoch, with 2s pre-stimulus period and 4s post-stimulus period. The variable baseline was corrected for in each trial by subtracting out the mean of the whole trial before doing further analyses.

To extract instantaneous phase information, single-trial data in each MEG channel were transformed using a Morlet wavelet function embedded in the Fieldtrip toolbox, with a frequency ranged from 1 to 60 Hz in steps of 1 Hz. To balance spectral and temporal resolution of the time-frequency transformation, from 1 to 20 Hz, the window length increased linearly from 1.5 cycles/frequency to 7 cycles/frequency, and was kept constant at 7 cycles/frequency above 20 Hz. The analysis windows for each trial was from 1s pre-stimulus period to 3s post-stimulus period with a temporal step of 10 ms. Phase and power response (squared absolute value) were extracted from the wavelet transform output at each time-frequency point for classification and mutual information analysis.

### Inter-trial phase coherence (ITPC)

In Experiment 1, the ‘inter-trial phase coherence’ (ITPC) was calculated on each time-frequency point (details as in (Lachaux et al. 1999)). ITPC is a measure of consistency of phase-locked neural activity entrained by stimuli across trials. ITPC of different frequency bands reflects phase tracking of cortical oscillations to temporally modulated stimuli. ITPC was computed across 20 trials for each of 3 TFS stimuli that were presented repeatedly and across 20 of 25 different TFS stimuli that were presented once each. As 1 – 5 trials were removed for each stimulus during preprocessing, to avoid bias of estimating ITPC caused by unequal numbers of trials across different stimuli, we only selected 20 trials for calculating ITPC.

### Single-trial classification

A single-trial classification analysis of 3 repeated TFS stimuli was carried out to investigate whether cortical oscillations entrained by TFS can differentiate the TFS from specific sentences. This classification analysis was described in detail in (Ng et al. 2013) as well as in (Luo and Poeppel 2007; Cogan and Poeppel 2011; Herrmann et al. 2013). For 20 trials of each repeated TFS stimulus, one trial was left out, and then a template was created by averaging phase series using the circular mean across the remaining trials for each of the TFS stimuli. There were three repeated TFS stimuli; so three templates of TFS stimuli were created. The circular distances between each template and each left-out trial from each TFS stimulus was computed. The circular distance was applied for phase classification by taking the circular mean during the period of 300 ms to 1500 ms after the stimulus onset, and on each frequency. A trial was given one template’s label if the distance between this trial and the template was the smallest among the three templates.

The classification analysis was conducted on each frequency within the frequency range that showed robust ITPC for 3 TFS stimuli compared with 25 TFS stimuli that were present once each.

A confusion matrix of classification scores was constructed for each trial of each stimulus type on each auditory channel. Then, classification performance was measured in a signal detection framework: correctly labelling the target stimulus was count as a ‘hit’ while labelling the other two stimuli as the target stimulus was counted as ‘false alarm’; *d*’ was calculated based on hit rates and false alarm rates and averaged across all auditory channels. Classification accuracy using the phase of each frequency was indicated by the mean of *d*’ over the three TFS stimuli, which was compared to the total *d*’ of the identification task which indicates participants’ sensitivity in the behavioral study (Macmillan and Creelman 2004)

### Spectral correlation and cochlear-scaled spectral correlation

The TFS preserves rich spectral information in speech, which may confer temporal information through the change of spectral content along time. To quantify this spectral change, inspired from cochlear-scaled entropy (Stilp et al. 2010), we created two indices – spectral correlation (SC) and cochlear-scaled spectral correlation (CSC). We first used a short-time Fourier transformation to generate spectral profiles of acoustic segments of 20 ms and then computed Pearson’s correlation coefficients of the spectral contents between adjacent temporal segments. This correlation, constrained by sampling rate and the number of samples in the sound files, is indicated as SC. Similar to the cochlear-scaled entropy, the spectral correlation computed after binning frequencies according to cochlear bands was indicated as CSC. We computed CSC using 32, 18, 8 and 4 bands separately to evaluate the effect of the number of cochlear bands on resolving temporal information from TFS.

### Recovered envelope from TFS

Previous studies have shown that speech amplitude envelopes can be recovered from TFS through cochlear processing (Ghitza 2001; Zeng et al. 2004). To measure how the recovered envelope from the TSF provides temporal information, we filtered 28 TFS stimuli in neurophysiological recording using Gammatone filterbanks of 32, 18, 8 and 4 bands, separately. The envelope of each cochlear band was extracted by using the Hilbert transform on each band and taking the absolute value (Glasberg and Moore 1990; Søndergaard and Majdak 2013). We then averaged the envelopes across all bands to get the recovered envelope from TFS.

### Mutual information (MI) analysis

To investigate what temporal information in TFS entrains neurophysiological responses and distinguishes different TFS sounds, we used the framework of MI to quantify shared information between MEG signals and acoustic properties in the stimuli (Quian Quiroga and Panzeri 2009; Panzeri et al. 2010). MI was calculated using the Information Breakdown Toolbox in MATLAB (Pola et al. 2003; Magri et al. 2009). We computed the MI between the phase series of each frequency (1 – 60 Hz) extracted from the time-frequency analysis described above and the acoustic properties of the stimuli in Experiment 1 - SC, CSC, original envelopes of sentences from which TFS stimuli were extracted, and the recovered envelopes from TFS (Cogan and Poeppel 2011; Gross et al. 2013; Ng et al. 2013; Kayser et al. 2015). The MI value of each frequency was calculated for each subject, for each condition, and for each auditory channel across trials.

In the present study, the acoustic properties of each stimulus are simply the values at each time point. For each frequency of the neurophysiological response, the phase distribution was composed of six equally spaced bins: 0 to pi/3, pi/3 to pi * 2/3, pi * 2/3 to pi, pi to pi * 4/3, pi * 4/3 to pi * 5/3, and pi * 5/3 to pi * 2. By choosing 6 bins for phase information, we ensured that there is enough temporal resolution to capture acoustic dynamics (Cogan and Poeppel 2011). The acoustic properties were first normalized within each sentence by dividing their maximum value and then grouped within each condition using 8 bins equally spaced from the minimum value to the maximum value. Eight bins were chosen because we wanted to have enough discrete precision to capture changes in acoustic properties while making sure that each bin has sufficient counts for MI analysis, since the greater number of bins would lead to zero counts in certain bins.

The estimation of MI is subject to bias caused by finite sampling of the probability distributions because limited data was supplied in the present study. Therefore, a quadratic extrapolation embedded in the Information Breakdown Toolbox was applied to correct bias. MI is computed on the data set of each condition. A quadratic function is then fit to the data points and the actual MI is taken to be the zero-crossing value. This new value reflects the estimated MI for an infinite number of trials and greatly reduces the finite sampling bias (Montemurro et al. 2007; Panzeri et al. 2007).

A shuffling procedure of sentence labels was applied to determine the baseline for MI analysis. 28 TFS sentences used in Experiment 1 were randomly assigned to trials in each condition to create a new date set and the same MI analysis was implemented on each frequency. This procedure was repeated 1000 times to get a distribution of MI values, from which a 99% one-side threshold was derived. By doing this, we preserved the structure of time series in data. Therefore, be comparing the baseline with the results, we ensured that MI results were not due to noise and specific processing procedures.

### Experiment 2: Behavioral measurement

All participants were sitting in a soundbooth while doing the tasks. In Experiment 2A, 10 R, 10 ER, and 10 FSR sentences for each window size were presented. We presented each type of sentence in separate blocks, such that three blocks (R block, ER block, and FSR block) were presented in Experiment 2A and there were 60 different R sentences (six window sizes × 10 sentences), 60 ER sentences (six window sizes × 10 sentences), and 30 FSR sentences (three window sizes × 10 sentences) total. Since we only have 100 different sentences in the materials, 10 sentences were shared between the R block and ER block. These shared ten sentences were in the largest window size condition (120 and 200 ms). Because intelligibility is very low for R and ER blocks under the largest window size, we can avoid the confound of hearing an intelligible sentence a second time which could prime listeners to better understand the second instance of the sentence.

Thirty different FSR sentences were selected from the sentences of the largest window size used in the R block and ER block, 15 from each block. The FSR block was always presented at the end, because, in our preliminary testing, we found that the FSR sentences were highly intelligible. We first presented the R block and then the ER block, instead of counter-balancing the order of two blocks. By doing this, we tried to avoid the confound that ER sentences contain more speech cues - as the intact TFS was preserved in ER sentences, participants could in theory adapt to the cues in ER sentences and improve performance in the R block.

In Experiment 2B, 10 ER and 10 ERNV sentences for each window size were presented and there were 60 ER sentences and 60 ERNV sentences total in two separate blocks. 10 sentences were shared between the ER block and the ERNV block. We set these ten shared sentences in the condition of the largest window size. The order of ER block and ERNV block was counter-balanced between participants.

Using MATLAB, the participants were presented with one sentence on each trial and were required to type in an Excel sheet what they heard after each sentence was presented (10 second limit). After the participants finished typing, they pushed a key on a message dialogue box to start next sentence. The input method for Chinese characters was Microsoft Pinyin IME 2003 without autocomplete. We treated each character of a sentence as one response and used the total number of the correct characters typed by the participants over 10 sentences divided by the total number of characters (70) as the intelligibility score for each window size.

### Psychometric function fitting

For the R and ER blocks in Experiment 2A and the ER block and ERNV block in Experiment 2B, we fitted a psychometric curve to each participant’s intelligibility score for each block using a Weibull function in the Palademes toolbox 1.5.2 (Prins and Kingdom 2009). The 50 percent intelligibility threshold and the slope of the psychometric curves were derived for each participant and later used for further analysis. Because intelligibility in FSR stayed at a ceiling level across all window sizes, we did not fit a psychometric curve to the data of FSR block. For the purpose of illustration, we averaged the intelligibility scores across subjects and fit a psychometric function to the averaged scores. These psychometric functions using group averaged scores were not used in any analysis.

## Results

### TFS robustly entrains low frequency cortical oscillatory responses

We used magnetoencephalography (MEG) recordings to measure neurophysiological responses to the TFS in Experiment 1. As cortical oscillatory responses entrained by auditory signals reflect how the auditory system both extracts temporal information and parses the incoming acoustic stream, studying cortical entrainment evoked by the TFS can reveal what information in TFS is extracted (Giraud and Poeppel 2012; Henry and Obleser 2012; Kayser et al. 2012; Luo and Poeppel 2012; Doelling et al. 2014; Henry et al. 2014; Kayser et al. 2015; Di Liberto et al. 2016; Teng et al. 2017).

We presented the 28 TFS stimuli in one block to 12 Mandarin Chinese native speakers while recording their neurophysiological responses. Three of the 28 TFS stimuli were repeatedly presented (25 times), and the remaining 25 TFS stimuli were presented once. We selected 20 MEG channels using a tone localizer (see Methods) and computed intertrial phase coherence (ITPC) across 20 trials for each of three repeated TFS stimuli and across TFS stimuli from 20 different sentences. We conducted a one-way repeated measures ANOVA on ITPC from 1 to 60 Hz, with four sentence types (three repeated sentences and one group of different sentences) as the main factor, and found a significant main effect from 1 – 8 Hz (*p* < 0.05, False Discovery Rate (FDR) corrected (Benjamini and Hochberg 1995)). We then grouped ITPC values from 1 to 8 Hz and conducted pairwise comparisons to examine differences between the four sentence types. The ITPC results show that cortical responses between 1 to 8 Hz are robustly entrained by the three repeated TFS stimuli (compared to the ITPC from the 20 distinct sentences; Fig 1B) (*p* < 0.05, paired t test, Bonferroni corrected). The topographies of ITPC show response patterns with an auditory origin, which is consistent with the hypothesis that the TFS evoked robust cortical entrainment (Fig. 1D).

To determine whether cortical oscillatory responses track distinct temporal structures reflected in the TFS, we employed a single-trial classifier to classify trials of the three repeated TFS stimuli between 1 and 8 Hz, using the MEG phase series (Fig. 1*C*) (Cogan and Poeppel 2011; Herrmann et al. 2013; Ng et al. 2013). We used a signal detection paradigm and converted the results of the classifier into the total *d*’, which indicates the classification accuracy across all of the three repeated TFS stimuli (See Methods). We found that the total *d*’ was significantly above the zero value (*d*’ for 33 percent correct classification) from 1 to 8 Hz (*p* < 0.05, one sample t test, Bonferroni corrected) (Fig. 1*C*, left panel). We then compared the total *d*’ between different frequencies and found that the phase patterns between 3 to 6 Hz were the most informative to the classification analysis (*p* < 0.05, paired t test, Bonferroni corrected).

These data on cortical entrainment to the TFS bear a strong resemblance to previous studies in which the SE was found to entrain neural responses below 10 Hz

(Luo and Poeppel 2007; Kerlin et al. 2010; Cogan and Poeppel 2011; Ding and Simon 2012; Peelle et al. 2013; Zion Golumbic et al. 2013). As prominent temporal information in speech between 3 and 6 Hz is carried by the SE (Ding et al. 2017), the results suggest that the temporal information relevant to the SE is perhaps read-out from the TFS by the auditory system.

### Temporal information extracted from TFS correlates with original envelope and can be reconstructed from spectral correlation

To investigate the nature of the temporal information extracted from the TFS, we measured how the SE could explain phase patterns evoked by the corresponding TFS using a mutual information (MI) framework (Quian Quiroga and Panzeri 2009; Panzeri et al. 2010). Analyses were conducted on the data from 11 of 12 Mandarin Chinese native speakers used in the previous analysis. One subject was excluded because of noise issues. We computed MI between the SE of 20 different sentences and the phase series evoked by the TFS derived from these 20 sentences. To set a baseline for MI values, we generated a null distribution of MI values by shuffling the labels between the sentences and computed MI values from 1000 shuffled datasets. We found that phase patterns of the entrained oscillations by TFS share a significant amount of information with the SE from 1 to 8 Hz (*p* < 0.01) (Fig. 2*A*, left panel), which aligns with the findings above on cortical entrainment (Fig. 1*A*). This result demonstrates that the temporal information extracted from the TFS strongly correlates with the envelope information.

Next we tried to determine how temporal information is extracted from the TFS. It has been suggested that the SE can be recovered from TFS through cochlear processing (Ghitza 2001; Zeng et al. 2004; Shamma and Lorenzi 2013). We filtered TFS stimuli using a gammatone filter bank of 16 bands to simulate cochlear processing (Patterson 1976; Patterson et al. 1987) and obtained the recovered envelopes from the gammatone filter outputs (See Methods for details). The averaged spectrum of recovered envelopes over the 25 sentences and an example of a time series can be seen in Figure 2*B* (red line). We then computed MI between the recovered envelope and the phase series evoked by the TFS. Although we found significant MI values across a wide range of frequencies (1 – 15 Hz) (*p* < 0.01, FDR corrected), the amount of MI between the recovered envelope and the phase series was much smaller than the MI between the SE and the phase series (Fig. 2*A*, red line). This result suggests that the recovered envelope is not solely what the auditory system uses to extract temporal information from TFS (Sheft et al. 2008). Other processes may be in play. Nonetheless, the result confirms the previous finding from a neurophysiological perspective, namely that when TFS is extracted using more than 8 bands, as in our present study, the recovered envelopes are no longer beneficial for speech recognition (Gilbert and Lorenzi 2006).

**Figure 2.**
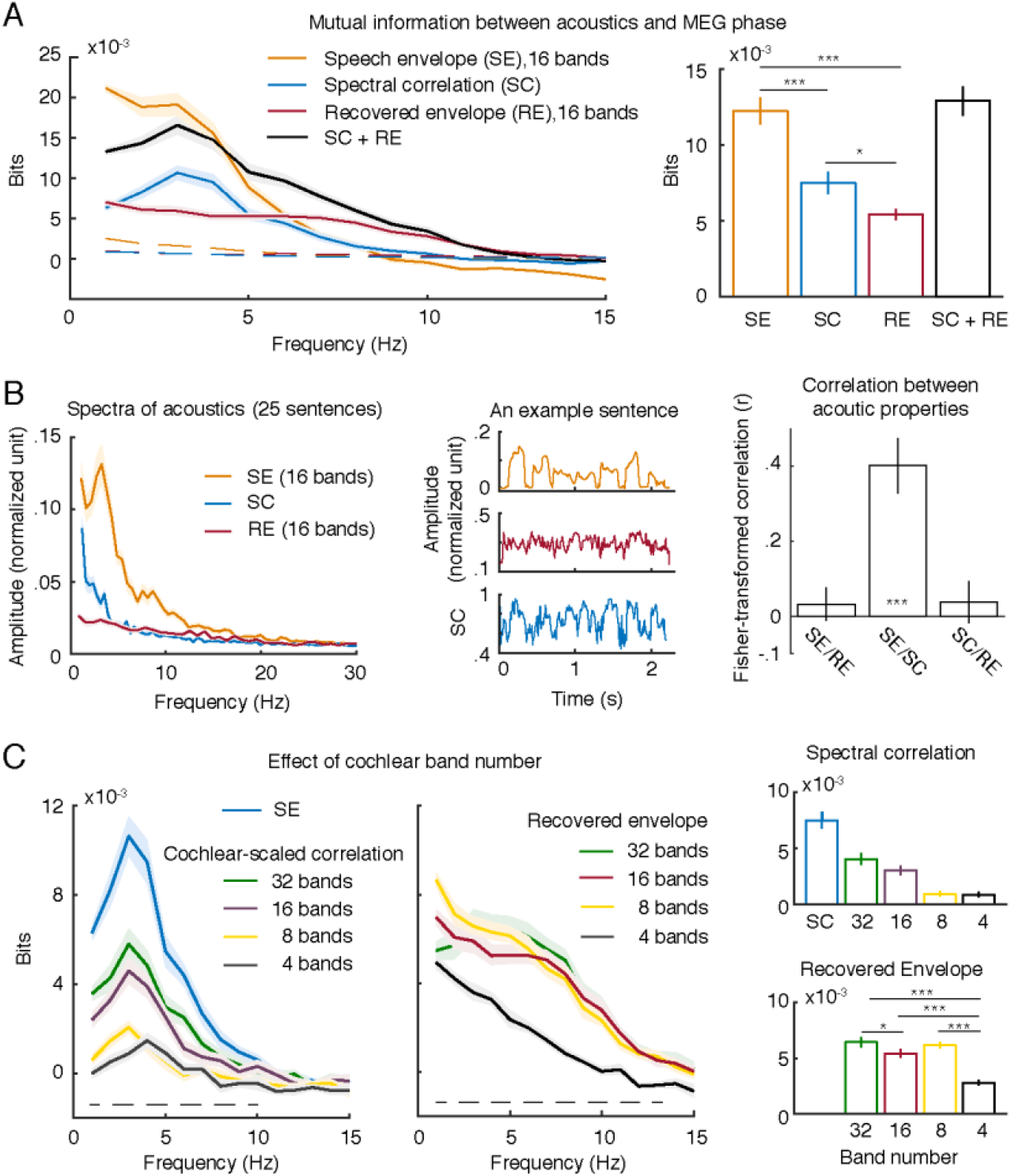
Results of mutual information (MI) analysis. (*A*) MI between MEG phase series evoked by the TFS and the speech amplitude envelope (SE), spectral correlation (SC), and recovered envelope (RE). The speech amplitude envelope and the recovered envelope were computed using 16 bands. The left panel shows MI results from 1 to 15 Hz. The dashed lines are significance thresholds with a one-sided alpha level of 0.01 for each acoustic property, which were derived from a permutation method (see Methods). The right panel shows average MI between 3 and 6 Hz where the phase series is most informative to discriminate the TFS from different sentences. The results show clearly that the TFS contains temporal information which correlates with the SE. The temporal information of the TFS can be derived from the spectral correlation and the recovered envelope. (*B*) Illustration of acoustics. The left panel shows the averaged spectra of the three acoustic properties used in the MI computation. The results were computed using the same 25 sentences for all the acoustic properties. The middle panel shows an example time series of each acoustic property for the same sentence. The color code is as in (*A*). The right panel shows Fisher-transformed Pearson correlation coefficients between three acoustic properties. It can be seen that the SE and the SC are highly correlated (*p* < 0.001). (C) Effects of spectral resolution on MI results. The left panel shows MI results of the spectral correlation and cochlear-scaled correlation. The dashed line indicates frequencies where the main effect of band number or spectral resolution is significant (*p* < 0.05, FDR corrected). The middle panel shows MI results of the recovered envelope computed using different number of frequency bands. The dashed line indicates frequencies where the main effect of band number is significant (*p* < 0.05, FDR corrected). The right panel shows averaged MI values between 3 and 6 Hz where the phase series is most informative. The results indicate that the spectral resolution or band number significantly affects the extraction of temporal information through spectral correlation - but not much through recovered envelope. This suggests that for cochlear implant users or people with hearing loss, the inability to use the TFS for speech perception could be due to degraded spectral resolution. The shaded area indicates +/- SEM over subjects. Asterisks show significance levels (***, *p* < 0.001; *, *p* < 0.05).

As TFS contains rich spectral information (Moore 2008; Shamma and Lorenzi 2013) and many studies have shown that cortical responses can be entrained by frequency modulations (Henry and Obleser 2012; Herrmann et al. 2013; Henry et al. 2014; Teng, Tian, Doelling, et al. 2017; Teng, Tian, Rowland, et al. 2017), the cortical entrainment elicited by the TFS may be caused by the spectral structure of the TFS. To reconstruct temporal information from the spectral structure of the TFS, we modified the cochlear-scaled entropy paradigm (Stilp and Kluender 2010) and computed spectral correlation using a short moving temporal window (20 ms) (see Methods for details). The average spectrum of spectral correlation of 25 sentences and an example of time series for one sentence can be seen in Figure 2*B* (blue line), which showed similar dynamics to the SE (Fig. 2*B*, orange line). The SE and the SC are significantly correlated over the 25 sentences used (one-sample t-test against zeros, *t*(24) = 12.16, *p* < 0.001, Bonferroni corrected), but no significant correlation was found between the RE and the SE (*t*(24) = 1.58, *p* = 0.384, Bonferroni corrected) as well as between the RE and the SC (*t*(24) = 1.45, *p* = 0.480, Bonferroni corrected) (Fig. 2*B*, right panel). We computed MI between the spectral correlation and the phase series evoked by TFS. MI values were found to be significant from 1 to 10 Hz (*p* < 0.01, FDR corrected) and were larger than the MI between the recovered envelope and the phase series from 2 to 4 Hz, which coincides with the peak of spectra of the SE (~ 3 Hz) (Fig. 2*A*, orange line). The results demonstrate that the auditory system extracts temporal information relevant to the SE from TFS using a procedure modelled by spectral correlation.

We combined MI computed from both spectral correlation and recovered envelope and summarized the MI values over 3 to 6 Hz, the most informative frequency range found in the classification analysis (Fig. 2*A*, right panel). We found that there was no significant difference between the MI values computed using SE and the MI values computed using both the recovered envelope and the spectral correlation (*t*(10) = 1.18, *p* = 1, *d* = 0.37, Bonferroni corrected).

In summary, the results suggest that temporal information can be extracted from TFS which correlates with the temporal information carried by the SE. A process that computes the spectral correlation between adjacent temporal frames could be used by the auditory system to extract the envelope information from TFS. The recovered envelope from cochlear processes also provides temporal information but does not seem to play a prominent role (Sheft et al. 2008).

### Extraction of temporal information from TFS depends on the number of frequency bands (spectral resolution)

As cochlear implants often provide poor spectral resolution (Oxenham and Kreft 2014) and persons with hearing loss manifest degraded spectral sensitivity (Hopkins and Moore 2011), we tested whether the number of frequency bands (spectral resolution) affects extracting temporal information from TFS, which may explain the inability of the cochlear implant users to effectively use TFS.

We computed the cochlear-scaled correlation (CSC) by binning frequencies into 32, 16, 8 and 4 cochlear bands separately (see Methods) and then calculated MI between MEG phase series and the CSC of the different band numbers (Fig. 2*C*, left panel). We conducted a one-way repeated measures ANOVA from 1 to 60 Hz, with spectral resolution as the main factor (five levels: SC, CSC of 32, 16, 8 and 4 bands), and found a significant main effect of the spectral resolution from 1 to 10 Hz (*p* < 0.05, FDR corrected). We then averaged MI from 3 to 6 Hz and found a downward linear trend with decreased spectral resolution (*F*(1,10) = 67.76, *p* < 0.001, η_p_^2^ = .871) (Fig. 2*C*, right panel). The results provide compelling evidence that spectral resolution significantly affects extraction of temporal information from the TFS.

We next examined the effect of the band number on the recovered envelope. We computed recovered envelopes using 32, 16, 8, and 4 cochlear bands separately and calculated MI between MEG phase series and the recovered envelopes of different band numbers (Fig. 2*C*, middle panel). We conducted a one-way repeated measures ANOVA from 1 to 60 Hz, with the number of cochlear bands as the main factor (four levels: 32, 16, 8 and 4 bands), and found a significant main effect of the band number from 1 to 13 Hz (*p* < 0.05, FDR corrected). We then averaged MI over 3 to 6 Hz and found a downward linear trend with decreased band number (*F*(1,10) = 61.93, *p* < 0.001, η_p_^2^ = .861). However, in a post-hoc test, we found that MI computed using 32 bands was larger than using 16 bands (*t*(10) = 4.18, *p* = .011, *d* = 1.32) but not more than 8 bands (*t*(10) = 0.80, *p* = 1, *d* = 0.25). The MI computed using 4 bands is lower than all the other bands (32 bands: *t*(10) = 9.89, *p* < .001, *d* = 3.13; 16 bands: *t*(10) = 6.86, *p* < .001, *d* = 2.17; 8 bands: *t*(10) = 10.89, *p* < .001, *d* = 3.44). Bonferroni correction was applied.

The results show that the number of bands does not affect recovering the envelope from the TFS as the frequency bands decrease from 32 to 8 bands. In contrast, we found a significant effect of the band number for spectral correlation. This is consistent with previous findings that spectral resolution modulates the efficiency of extracting temporal information from the TFS (Léger et al. 2015; Oxenham 2018).

### TFS significantly compensates for SE temporal information compromised in temporally disrupted speech

We tested whether the TFS compensates for lost temporal information when the SE is disrupted in Experiment 2. We used a ‘reversed speech’ paradigm in which the temporal structure of speech signals is compromised by temporally reversing local segments and then tested speech intelligibility (Saberi and Perrott 1999; Stilp et al. 2010). The rationale is that the reversing procedure disrupts the modulation phase of the SE and, as the reversed temporal window becomes larger, the modulation phase gets distorted more severely and provides incorrect cues on the temporal structure of speech. This reversing procedure, therefore, renders the temporal cues carried by the reversed SE unavailable for segmenting speech signals. Then we can test whether the auditory system extracts certain cues provided by the TFS to segment speech signals. Figure 3*A* show a schematic illustration of stimulus generation. By comparing speech intelligibility for the reversed speech when TFS is intact with the conditions when the SE and TFS are both disrupted, we determine whether the TFS, similar to the SE, provides critical temporal information for speech segmentation.

**Figure 3.**
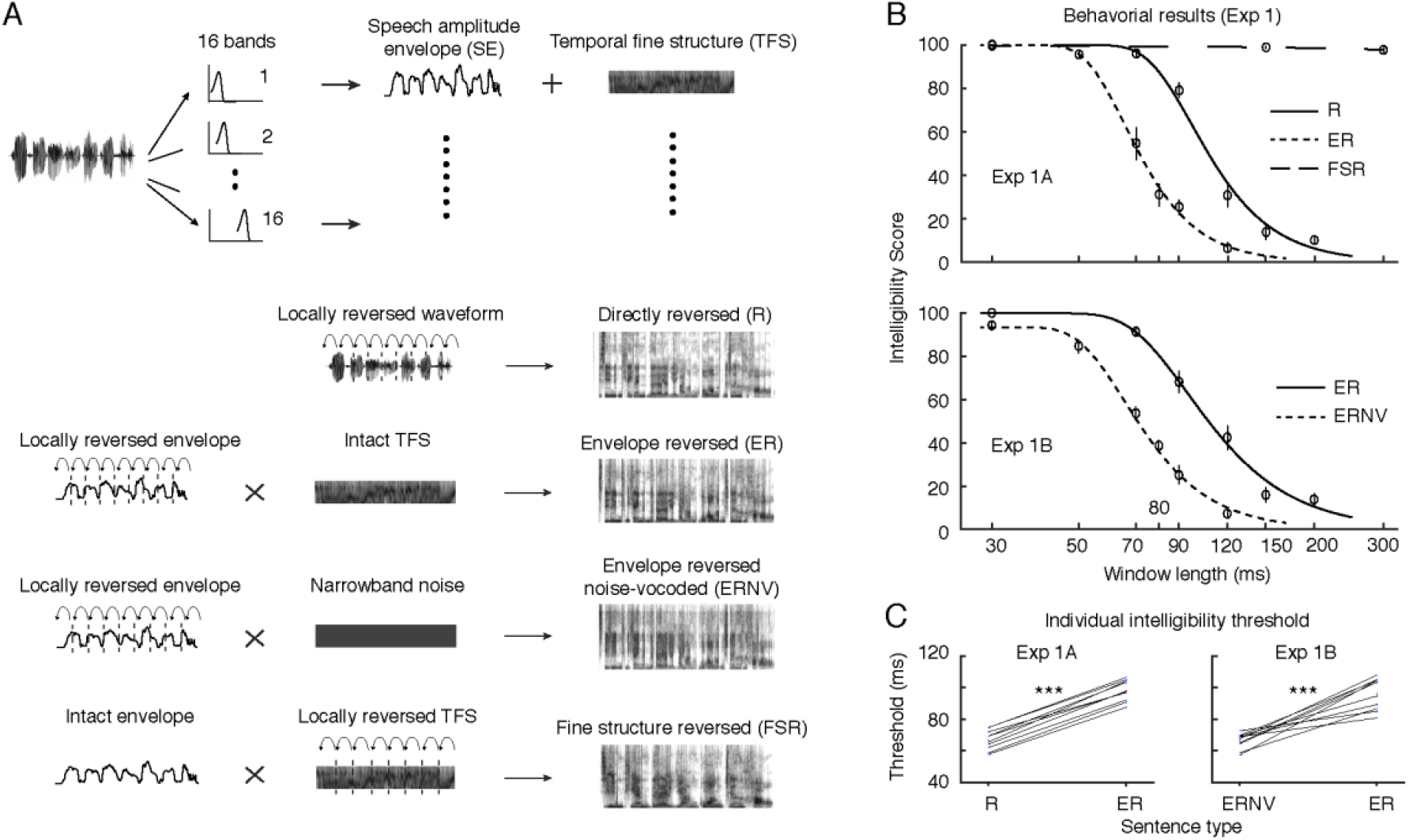
Illustration of stimulus manipulations and behavioral results. (A) Speech signals were first decomposed into the amplitude envelope (SE) and the temporal fine structure (TFS). Rectangular windows with various lengths were used to segment speech signals and then locally reverse each speech segment in time. Depending on which part of the speech signal was locally reversed, four types of reversed sentences were generated: directly reversed (R) – raw broadband speech signals were locally reversed; envelope reversed (ER) – we reversed only the envelope and then used it to modulate the intact TFS; envelope reversed noise-vocoded (ERNV) – we reversed the envelope and used it to modulate narrow band noise; fine structure reversed (FSR) – we reversed the TFS and used the intact envelope to modulate the reversed TFS. (B) Behavioral results of Intelligibility of Experiments 2A and 2B. The top panels show group-averaged results of Experiment 2A and Experiment 2B and the psychometric functions fit to the data. The x-axis represents window length, the y-axis represents the intelligibility score. It can be clearly seen that ER sentences with intact TFS show significantly higher intelligibility than sentences without intact TFS (R in Expt. 2A and ERNV in Expt. 2B). The lower panel shows 50% thresholds of psychometric function fits to individual data. Each black line represents each subject’s individual threshold. The x-axis shows sentence type and the y-axis the threshold. Asterisks show significance levels (***, *p* < 0.001). The error bars are +/- SEM over subjects.

In Experiment 2A, we directly reversed the speech segments and varied the segment size to generate directly reversed (R) sentences – i.e. SE and TFS were both reversed. Next, we locally reversed the SE and used this reversed SE to modulate the intact TFS to generate envelope reversed (ER) sentences. For fine structure reversed (FSR) sentences, we used the intact SE to modulate the reversed TFS. An illustration of the stimulus generation and spectrograms of one example sentence is shown in Fig 3A (See Methods for details). We recruited 10 Chinese native speakers and tested speech intelligibility for these sentences (ten sentences for each segment size). For Experiment 1B, we created envelope reversed noise-vocoded (ERNV) sentences by using the reversed SE to modulate noise (Fig 3A). By comparing intelligibility of 10 Chinese native speakers for the ER sentences with the ERNV sentences, we further evaluated the contribution of the TFS. The behavioral results are shown in Fig 3B.

We fit each participant’s intelligibility scores to a psychometric function (see Methods) and conducted a two-way mixed effect ANOVA on the thresholds of the psychometric functions (fifty percent correct threshold). We treated Experiment (Experiment 2A and Experiment 2B) as the between-subjects factor and TFS (with intact TFS: ER, or without intact TFS: R and ERNV) as the within-subject factor. We found a significant main effect of TFS (*F*(1,18) = 227.14, *p* < 0.001, η_p_^2^ = .927), but not for Experiment (*F*(1,18) = .233, *p* = 0.635, η_p_^2^ = .013). In a post-hoc test, we found in Experiment 1A that thresholds (which are best interpreted as temporal tolerance with respect to the reversal distortion) in the ER block were significantly larger than in the R block (paired sample t-test: *t*(9) = 26.41, *p* < .001, *d* = 8.35, 95% CI [33.83, 29.49]); in Experiment 1B, thresholds in the ER block were significantly larger than the ERNV block (paired sample t-test: *t*(9) = 7.71, *p* < .001, *d* = 2.44, 95% CI [38.65, 21.12]).

The behavioral results for R sentences replicate previous findings that the intelligibility of locally reversed speech degrades with increased segment size (Saberi and Perrott 1999; Kiss et al. 2008; Stilp et al. 2010). Although Mandarin is a tone language, and one might have expected tone-related differences in behavioral thresholds, the results of R sentences are comparable to the results of previous studies using sentences from non-tonal languages (Saberi and Perrott 1999; Stilp et al. 2010).

Next we turn to the novel manipulations. In Experiment 2A, speech intelligibility for the ER sentences was significantly higher than for the R sentences. As the key difference between the R and ER sentences is only whether the TFS is disrupted, this result suggests that the intact TFS in the ER sentences aids in restoring the temporal information in the reversed speech by increasing intelligibility. In Experiment 2B, we also found that the intelligibility threshold for the ER sentences was significantly larger than for the ERNV sentences, and that the intelligibility difference between ERNV and ER sentences was comparable to the difference between the R and ER sentences. This second result lends strong support to the hypothesis that the auditory system extracts temporal information from the TFS to compensate for the disrupted SE.

Speech intelligibility for the FSR sentences did not change across different segment sizes. This could be because the intact SE supplied sufficient information for speech segmentation even though TFS was compromised. This could be considered a similar case as Shannon et al. (1995), where noise modulated by four bands of the SE was sufficient for speech perception. The data invite the hypothesis that the temporal information in the TFS only starts to become a significant cue when the SE is disrupted.

The gain in speech intelligibility by adding the intact TFS may be caused by an interaction at the acoustic level between TFS and the reversed envelope, such that the original envelope is recovered (Kates 2011; Shamma and Lorenzi 2013). Admittedly, while the shape of the envelope can be changed depending on its carrier, we argue that this contributes little to the gain in intelligibility. In Figure 4, we show that the spectra of the envelopes of the ER and ERNV sentences do not significantly differ across different segment sizes. We first generated the ER and ERNV sentences from 20 raw sentences and then extracted their envelopes using 16 bands. We computed the average spectra of the ER and ERNV envelopes (Fig. 4*A*, upper panel) and then the differential spectra between the ER and ERNV envelopes (Fig. 4*A*, lower panel). To quantify whether the differences between the ER and ERNV envelope spectra were significant, we computed a bootstrap threshold with a two-sided alpha level of 0.05: we randomly sampled from the ER and ERNV sentences to form two new groups of sentences and then computed a differential spectrum. We repeated this procedure 1000 times to generate thresholds for the differential spectra between ER and ERNV. For window lengths of 70 and 90 ms, speech intelligibility differed significantly between ER and ERNV sentences (Fig. 3*C*), but there were no significant differences of the spectra of the envelopes (Fig. *4B*).

**Figure 4.**
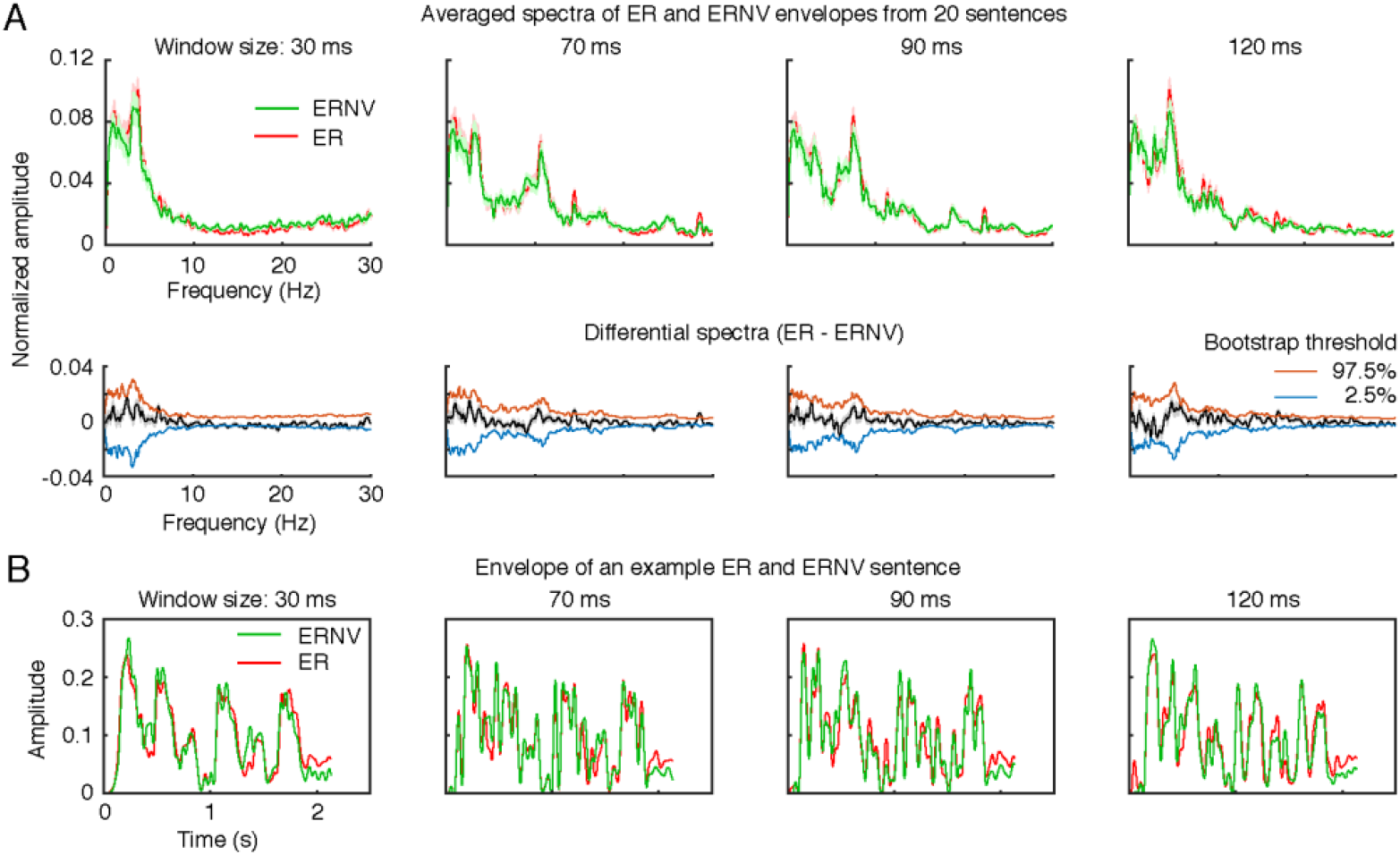
(A) The upper panels: averaged spectra of the envelopes from 20 ER and ERNV sentences across four window lengths used in Experiment 1. The green line represents the ERNV sentences; the red line represents the ER sentences. The x-axis is frequency and the y-axis shows normalized amplitude of the spectra. The lower panel depicts the differential spectra computed by subtracting the spectra of ERNV sentences from the spectra of ER sentences. The black line represents the differential spectra and the red and blue lines show bootstrap thresholds. The envelopes of the ER and ERNV sentences do not significantly differ. Note that this is also true for the window lengths of 70 ms and 90 ms, where speech intelligibility for the ER sentences was significantly higher than the ERNV sentences. (B) Example envelopes of an ER and ERNV sentence across different window lengths. The shaded area represents +/- SEM over sentences.

## Discussion

Previous studies that principally focused on the speech amplitude envelope revealed only part of the mechanism for segmentation. The SE and the TFS convey comparable temporal information to elicit cortical entrainment. The auditory system relies on various cues in speech signals, both temporal and spectral, for segmentation. The data we presented demonstrate that the TFS contains temporal information that can be used for speech segmentation by entraining cortical oscillatory responses. Using MEG experiments, we first demonstrated that the TFS could entrain cortical oscillatory responses, with phase patterns specific to a particular TFS. We then evaluated contributions to temporal information from the recovered envelope and the spectral dynamics in the TFS and found that the temporal information of the TFS comes primarily from the dynamics of its spectral structure, which can be captured using spectral correlation. A further analysis on the effect of the number of frequency bands showed that spectral resolution plays a major role in extracting temporal information from TFS. Using behavioral measurements, we next showed that the TFS contains critical acoustic cues for the auditory system to restore the temporal structure of speech and aids speech recognition.

### The data explain previous findings on the benefit of TFS in challenging listening environments

Our behavioral results showed that TFS helps the auditory system restore temporal information when the SE is disrupted. This result echoes previous findings that TFS helps increase speech intelligibility under challenging environments (Hopkins et al. 2008; Moore 2008) and further suggests that the gain from the TFS is a result of complementary temporal information contained in the TFS. The auditory system can monitor spectral changes to recover temporal information lost due to a disrupted envelope.

Our results involving cortical entrainment evoked by the TFS could explain the finding that, when the SE is smoothed by added noise, robust cortical entrainment can still be found (Zoefel and VanRullen 2015). The smoothing procedure used in Zoefel et al. (2015) operated on each frequency band and could have left the TFS intact and therefore, the auditory system could still extract sufficient temporal information from the spectral structure of the processed speech signals. Other data show that the TFS helps the speech amplitude envelope entrain cortical oscillatory responses in noise (Ding et al. 2014), which is presumably because the TFS contains temporal information similar to the SE and can provide comparable temporal information to entrain cortical oscillatory responses when the envelope is disrupted by white noise.

### The auditory system extracts temporal information from both the TFS and the SE

Previous studies (Ghitza 2001; Zeng et al. 2004) suggest that the auditory system extracts temporal information from the TFS through the recovered envelope. Our data support this general point. Furthermore, we also found that the recovered envelope by itself cannot fully explain the amount of temporal information in the TFS (Fig. 2*A*). Our MI analysis between SC and the MEG phase series reveals that the auditory system monitors spectral changes in the TFS to extract temporal information, and that the dynamics of the spectral structure in the TFS explains a larger amount of variance, indicated by MI analysis, in the MEG phase series than the recovered envelope.

The TFS preserves the rich spectral information of speech and carries information relevant to pitch (Moore and Moore 2003; Moore et al. 2006), lexical tone (Xu and Pfingst 2003; Zeng et al. 2005), and the acoustic transitions between consonants and vowels (Rosen 1992). All this information in the spectral domain of speech provides dynamic cues for speech segmentation. We used SC and CSC to extract temporal information from TFS, which we aim to reveal in the TFS how spectral information changes along time and where temporal information exists. This does not necessarily mean that SC and CSC represent how exactly the auditory system extracts information from the TFS. SC and CSC can be viewed as one of many algorithms that the auditory system implements to process the TFS, and other algorithms may also achieve a similar computational goal of recovering temporal information from the TFS (Shamma and Lorenzi 2013; Ewert et al. 2018).

### Spectral resolution plays a key role in processing the TFS

We found that the number of frequency bands strongly modulates the extraction of temporal information from the TFS (Fig. 2*C*). Although temporal acuity, which is important for processing temporal fluctuations in speech, is relatively preserved in cochlear implant users, deficits of perceiving speech in challenging environments among people with hearing loss still exists (Oxenham and Kreft 2014). It has been argued that the reduced spectral resolution smears spectral details of speech and maskers (Fu and Nogaki 2005; Oxenham and Kreft 2014). Our results suggest that the reduced spectral resolution smears the spectral details of TFS which results in poor extraction of temporal information from the spectral structure in speech. This smearing effect prevents the auditory system from parsing the speech stream using spectral contents and therefore leads to degraded intelligibility.

The correlation between spectral resolution and speech intelligibility (Oxenham and Kreft 2014) could be due to the way the auditory system groups speech information in terms of spectral and temporal structure. Modulated noise often reduces listeners’ sensitivity to the TFS (Hopkins and Moore 2011), which could be because modulated noise contains dynamic changes in spectral contents and interferes with the spectral structure preserved in TFS. Reduced spectral resolution prevents the auditory system both from tracking spectral changes in speech signals and from separating maskers from target speech based on spectral details.

### The relationship between the envelope and fine structure

Speech is often investigated as an envelope and a fine structure. As the envelope manifests characteristic temporal structures of speech signals, such as syllable and phoneme structures (Di Liberto et al. 2016), mechanisms revealed by studies on the envelope are often argued to be related to temporal coding. Therefore, many studies on speech segmentation mostly focus on the SE and separate between the SE and the TFS. The present study indicates a blurred line between the envelope and TFS in speech segmentation, as our results show that the auditory system can rely on the TFS to temporally group speech information. The common practice of using band-filtering methods to separate envelope and TFS may not be an ideal way to extract critical features from speech signals, as previous finding shows that the TFS and the SE code comparable information in the early/periphery auditory system and therefore cannot be fully separated (Shamma and Lorenzi 2013). Speech segmentation could rely on local temporal-spectral operations, such as SC and CSC used in the present study and spectro-temporal receptive fields found in many studies (Theunissen et al. 2000; Eggermont 2001; Machens et al. 2004; Ding and Simon 2013; Mesgarani et al. 2014).

We calculated the modulation power spectra of raw speech signals and the TFS, using different numbers of frequency bands (Singh and Theunissen 2003; Elliott and Theunissen 2009). We created time-frequency representations of the speech signals and TFS using the log amplitude of their spectrograms obtained with Gaussian windows. We then applied the 2D Fourier Transform to the spectrograms and created modulation power spectra by taking the amplitude squared as a function of the Fourier pairs of the time and frequency axes. We illustrate in Figure 5 that the TFS represents the residual signals left by envelope extraction - the higher the number of frequency bands used, the less modulation power was left in the TFS. This demonstration further argues that TFS and envelope cannot be fully separated and the auditory system segments speech signals in a ‘holistic’ or ‘synthetic’ manner, concurrently and cooperatively using both temporal and spectral information.

**Figure 5.**
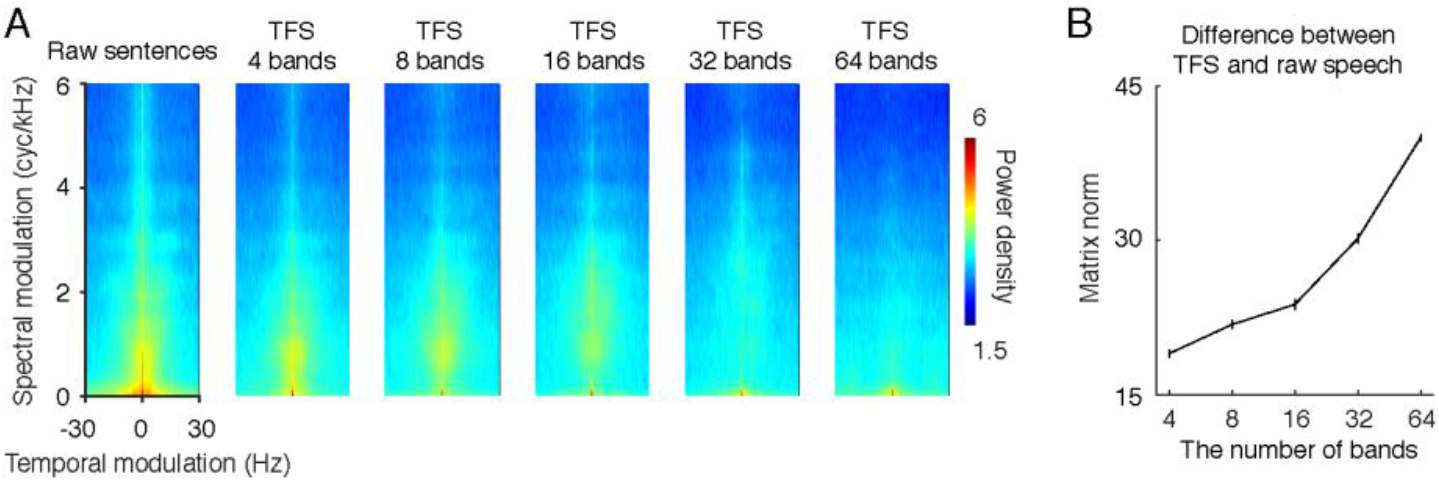
Averaged modulation power spectrum of speech signals and TFS. (*A*)We selected 20 sentences used in the present study for analysis and averaged the results across 20 sentences. The x-axis of each plot represents temporal modulation and the y-axis spectral modulation. From left to right, the modulation power spectrum was calculated for raw sentences and the TFS reconstructed using different numbers of frequency bands. The modulation power decreases as more frequency bands are used to extract the envelope. (*B*) We quantified the difference of the modulation power spectrum between the raw sentences and the TFS by first calculating subtraction between the modulation power spectrum of the raw sentences and the TFS and then taking the norm. It can be seen that, as the number of frequency bands increases, the TFS preserves less modulation information of the raw sentences (the matrix norm increases).

### Conclusion: speech segmentation from a synthetic perspective

The envelope and the temporal fine structure of speech are often studied separately in the context of cortical entrainment to speech and speech segmentation. Our study demonstrates that the auditory system treats speech as a whole, and that temporal speech segmentation involves processing both the temporal and spectral contents of speech. This view provides new interpretations to previous findings and also generates new hypotheses for the future study of the neural basis of speech segmentation.

## Acknowledgements

We thank Jeff Walker for his technical support and Ning Mei for his assistance on collecting and organizing data. This research was supported by NIH 5R01DC05660 to DP and the Max-Planck-Society.

Author Contributions
Conceived and designed the experiments: XT, GBC & DP
Performed the experiments: XT
Analyzed the data: XT
Contributed reagents/materials/analysis tools: XT
Wrote the paper: XT, GBC, DP

## References

Benjamini Y, Hochberg Y. 1995. Controlling the false discovery rate: A practical and powerful approach to multiple testing. J Roy Stat Soc B Met. 57:289–300.

Cogan GB, Poeppel D. 2011. A mutual information analysis of neural coding of speech by low-frequency MEG phase information. J Neurophysiol. 106:554–563.

de Cheveigné A, Simon JZ. 2007. Denoising based on time-shift PCA. J Neurosci Meth. 165:297–305.

de Cheveigné A, Simon JZ. 2008. Sensor noise suppression. J Neurosci Meth. 168:195–202.

Di Liberto GM, O’Sullivan JA, Lalor EC. 2016. Low-Frequency Cortical Entrainment to Speech Reflects Phoneme-Level Processing. Curr Biol. 25:2457–2465.

Ding N, Chatterjee M, Simon JZ. 2014. Robust cortical entrainment to the speech envelope relies on the spectro-temporal fine structure. NeuroImage. 88 IS -:41–46.

Ding N, Patel AD, Chen L, Butler H, Luo C, Poeppel D. 2017. Temporal modulations in speech and music. Neurosci Biobehav R. 81:181–187.

Ding N, Simon JZ. 2012. Emergence of neural encoding of auditory objects while listening to competing speakers. Proc Natl Acad Sci U S A. 109:11854–11859.

Ding N, Simon JZ. 2013. Adaptive Temporal Encoding Leads to a Background-Insensitive Cortical Representation of Speech. J Neurosci. 33:5728–5735.

Ding N, Simon JZ. 2014. Cortical entrainment to continuous speech: functional roles and interpretations. Front Hum Neurosci. 8:1–7.

Doelling KB, Arnal LH, Ghitza O, Poeppel D. 2014. Acoustic landmarks drive delta-theta oscillations to enable speech comprehension by facilitating perceptual parsing. NeuroImage. 85, Part 2 IS -:761–768.

Eggermont JJ. 2001. Temporal modulation transfer functions in cat primary auditory cortex: separating stimulus effects from neural mechanisms. J Neurophysiol. 87:305–321.

Elliott TM, Theunissen FE. 2009. The Modulation Transfer Function for Speech Intelligibility. PLoS Comput Biol. 5:1–14.

Ewert SD, Paraouty N, Lorenzi C. 2018. A two-path model of auditory modulation detection using temporal fine-structure and envelope cues. European Journal of Neuroscience.

Fu Q-J, Nogaki G. 2005. Noise Susceptibility of Cochlear Implant Users: The Role of Spectral Resolution and Smearing. JARO. 6:19–27.

Fu Q-J, Zhu M, Wang X. 2011. Development and validation of the Mandarin speech perception test. J Acoust Soc Am. 129:EL267–EL273.

Ghitza O. 2001. On the upper cutoff frequency of the auditory critical-band envelope detectors in the context of speech perception. J Acoust Soc Am. 110:1628–1640.

Ghitza O. 2012. On the Role of Theta-Driven Syllabic Parsing in Decoding Speech: Intelligibility of Speech with a Manipulated Modulation Spectrum. Front Psychol. 3.

Ghitza O, Greenberg S. 2009. On the Possible Role of Brain Rhythms in Speech Perception: Intelligibility of Time-Compressed Speech with Periodic and Aperiodic Insertions of Silence. Phonetica. 66:113–126.

Gilbert G, Bergeras I, Voillery D, Lorenzi C. 2007. Effects of periodic interruptions on the intelligibility of speech based on temporal fine-structure or envelope cues. J Acoust Soc Am. 122:1336–1339.

Gilbert G, Lorenzi C. 2006. The ability of listeners to use recovered envelope cues from speech fine structure. J Acoust Soc Am. 119:2438–2444.

Giraud A-L, Poeppel D. 2012. Cortical oscillations and speech processing: emerging computational principles and operations. Nat Neurosci. 15:511–517.

Glasberg BR, Moore BCJ. 1990. Derivation of auditory filter shapes from notched-noise data. Hear Res. 47:103–138.

Gross J, Hoogenboom N, Thut G, Schyns P, Panzeri S, Belin P, Garrod S. 2013. Speech rhythms and multiplexed oscillatory sensory coding in the human brain. PLoS Biol. 11:e1001752.

Henry MJ, Herrmann B, Obleser J. 2014. Entrained neural oscillations in multiple frequency bands comodulate behavior. Proc Natl Acad Sci U S A. 111:14935–14940.

Henry MJ, Obleser J. 2012. Frequency modulation entrains slow neural oscillations and optimizes human listening behavior. Proc Natl Acad Sci U S A. 109:20095–20100.

Herrmann B, Henry MJ, Grigutsch M, Obleser J. 2013. Oscillatory Phase Dynamics in Neural Entrainment Underpin Illusory Percepts of Time. J Neurosci. 33:15799–15809.

Hopkins K, Moore BCJ. 2011. The effects of age and cochlear hearing loss on temporal fine structure sensitivity, frequency selectivity, and speech reception in noise. J Acoust Soc Am. 130:334–349.

Hopkins K, Moore BCJ, Stone MA. 2008. Effects of moderate cochlear hearing loss on the ability to benefit from temporal fine structure information in speech. J Acoust Soc Am. 123:1140–1153.

Hopkins K, Moore BCJ, Stone MA. 2010. The effects of the addition of low-level, low-noise noise on the intelligibility of sentences processed to remove temporal envelope information. J Acoust Soc Am. 128:2150–2161.

Kates JM. 2011. Spectro-temporal envelope changes caused by temporal fine structure modification. J Acoust Soc Am. 129:3981–3990.

Kayser C, Ince RAA, Panzeri S. 2012. Analysis of slow (theta) oscillations as a potential temporal reference frame for information coding in sensory cortices. PLoS Comput Biol. 8:e1002717.

Kayser SJ, Ince RAA, Gross J, Kayser C. 2015. Irregular Speech Rate Dissociates Auditory Cortical Entrainment, Evoked Responses, and Frontal Alpha. J Neurosci. 35:14691–14701.

Kerlin JR, Shahin AJ, Miller LM. 2010. Attentional Gain Control of Ongoing Cortical Speech Representations in a “Cocktail Party.” J Neurosci. 30:620–628.

Kiss M, Cristescu T, Fink M, Wittmann M. 2008. Auditory language comprehension of temporally reversed speech signals in native and non-native speakers. Acta neurobiologiae experimentalis. 68:204.

Lachaux J-P, Rodriguez E, Martinerie J, Varela FJ. 1999. Measuring phase synchrony in brain signals. Hum Brain Mapp. 8:194–208.

Léger AC, Reed CM, Desloge JG, Swaminathan J, Braida LD. 2015. Consonant identification in noise using Hilbert-transform temporal fine-structure speech and recovered-envelope speech for listeners with normal and impaired hearinga). J Acoust Soc Am. 138:389–403.

Lorenzi C, Gilbert G, Carn H, Garnier S, Moore BCJ. 2006. Speech perception problems of the hearing impaired reflect inability to use temporal fine structure. Proc Natl Acad Sci U S A. 103:18866–18869.

Luo H, Poeppel D. 2007. Phase patterns of neuronal responses reliably discriminate speech in human auditory cortex. Neuron. 54:1001–1010.

Luo H, Poeppel D. 2012. Cortical oscillations in auditory perception and speech: evidence for two temporal windows in human auditory cortex. Front Psychol. 3:170.

Machens CK, Wehr MS, Zador AM. 2004. Linearity of Cortical Receptive Fields Measured with Natural Sounds. J Neurosci. 24:1089–1100.

Macmillan NA, Creelman CD. 2004. Detection Theory: A User’s Guide. Taylor & Francis.

Magri C, Whittingstall K, Singh V, Logothetis NK, Panzeri S. 2009. A toolbox for the fast information analysis of multiple-site LFP, EEG and spike train recordings. BMC Neurosci. 10:81.

Mesgarani N, Cheung C, Johnson K, Chang EF. 2014. Phonetic Feature Encoding in Human Superior Temporal Gyrus. Science. 343:1006–1010.

Montemurro MA, Senatore R, Panzeri S. 2007. Tight data-robust bounds to mutual information combining shuffling and model selection techniques. Neural Comput. 19:2913–2957.

Moore BCJ. 2008. The Role of Temporal Fine Structure Processing in Pitch Perception, Masking, and Speech Perception for Normal-Hearing and Hearing-Impaired People. JARO. 9:399–406.

Moore BCJ, Glasberg BR, Flanagan HJ, Adams J. 2006. Frequency discrimination of complex tones; assessing the role of component resolvability and temporal fine structure. J Acoust Soc Am. 119:480–490.

Moore GA, Moore BCJ. 2003. Perception of the low pitch of frequency-shifted complexes. J Acoust Soc Am. 113:977–985.

Ng BSW, Logothetis NK, Kayser C. 2013. EEG phase patterns reflect the selectivity of neural firing. Cereb Cortex. 23:389–398.

Oldfield RC. 1971. The assessment and analysis of handedness: The Edinburgh inventory. Neuropsychologia. 9:97–113.

Oostenveld R, Fries P, Maris E, Schoffelen J-M. 2011. Fieldtrip: open source software for advanced analysis of MEG, EEG, and invasive electrophysiological data. Comput Intell Neurosci. 2011:1–9.

Oxenham AJ. 2018. How We Hear: The Perception and Neural Coding of Sound. Annu Rev Psychol. 69:27–50.

Oxenham AJ, Kreft HA. 2014. Speech Perception in Tones and Noise via Cochlear Implants Reveals Influence of Spectral Resolution on Temporal Processing. Trends in Hearing. 18:2331216514553783.

Panzeri S, Brunel N, Logothetis NK, Kayser C. 2010. Sensory neural codes using multiplexed temporal scales. Trends Neurosci. 33:111–120.

Panzeri S, Senatore R, Montemurro MA, Petersen RS. 2007. Correcting for the Sampling Bias Problem in Spike Train Information Measures. J Neurophysiol. 98:1064–1072.

Patterson RD. 1976. Auditory filter shapes derived with noise stimuli. J Acoust Soc Am. 59:640.

Patterson RD, Nimmo-Smith I, Holdsworth J, Rice P. 1987. An efficient auditory filterbank based on the gammatone function. Ina meeting of the IOC Speech Group on Auditory Modelling at RSRE. 2.

Peelle JE, Gross J, Davis MH. 2013. Phase-locked responses to speech in human auditory cortex are enhanced during comprehension. Cerebral Cortex. 23:1378–1387.

Poeppel D. 2003. The analysis of speech in different temporal integration windows: cerebral lateralization as “asymmetric sampling in time.” Speech Commun. 41:245–255.

Pola G, Thiele A, Hoffmann KP, Panzeri S. 2003. An exact method to quantify the information transmitted by different mechanisms of correlational coding. Network. 14:35–60.

Prins N, Kingdom FAA. 2009. Palamedes: Matlab routines for analyzing psychophysical data. http://www.palamedestoolbox.org.

Quian Quiroga R, Panzeri S. 2009. Extracting information from neuronal populations: information theory and decoding approaches. Nat Rev Neurosci. 10:173–185.

Roberts TP, Ferrari P, Stufflebeam SM, Poeppel D. 2000. Latency of the auditory evoked neuromagnetic field components: stimulus dependence and insights toward perception. J Clin Neurophysiol. 17:114–129.

Rosen S. 1992. Temporal Information in Speech: Acoustic, Auditory and Linguistic Aspects. Philosophical Transactions of the Royal Society of London B: Biological Sciences. 336:367–373.

Saberi K, Perrott DR. 1999. Cognitive restoration of reversed speech : Abstract : Nature. Nature. 398:760–760.

Shamma S, Lorenzi C. 2013. On the balance of envelope and temporal fine structure in the encoding of speech in the early auditory system. J Acoust Soc Am. 133:2818–2833.

Shannon RV, Zeng FG, Kamath V, Wygonski J. 1995. Speech Recognition with Primarily Temporal Cues. Science.

Sheft S, Ardoint M, Lorenzi C. 2008. Speech identification based on temporal fine structure cues. J Acoust Soc Am. 124:562–575.

Singh NC, Theunissen FE. 2003. Modulation spectra of natural sounds and ethological theories of auditory processing. J Acoust Soc Am. 114:3394–3411.

Smith ZM, Delgutte B, Oxenham AJ. 2002. Chimaeric sounds reveal dichotomies in auditory perception. Nature. 416:87–90.

Stilp CE, Kiefte M, Alexander JM, Kluender KR. 2010. Cochlea-scaled spectral entropy predicts rate-invariant intelligibility of temporally distorted sentences. J Acoust Soc Am. 128:2112–2126.

Stilp CE, Kluender KR. 2010. Cochlea-scaled entropy, not consonants, vowels, or time, best predicts speech intelligibility. Proc Natl Acad Sci U S A. 107:12387–12392.

Swaminathan J, Mason CR, Streeter TM, Best V, Roverud E, Kidd G. 2016. Role of Binaural Temporal Fine Structure and Envelope Cues in Cocktail-Party Listening. J Neurosci. 36:8250–8257.

Søndergaard PL, Majdak P. 2013. The Auditory Modeling Toolbox. In: Blauert J editor. The Technology of Binaural Listening. Berlin, Heidelberg: Springer Berlin Heidelberg. p. 33–56.

Teng X, Tian X, Rowland J, Poeppel D. 2017. Concurrent temporal channels for auditory processing: Oscillatory neural entrainment reveals segregation of function at different scales. PLoS Biol. 15:e2000812.

Theunissen FE, Sen K, Doupe AJ. 2000. Spectral-temporal receptive fields of nonlinear auditory neurons obtained using natural sounds. J Neurosci. 20:2315–2331.

Xu L, Pfingst BE. 2003. Relative importance of temporal envelope and fine structure in lexical-tone perception (L). J Acoust Soc Am. 114:3024–3027.

Zeng F-G, Nie K, Liu S, Stickney G, Del Rio E, Kong Y-Y, Chen H. 2004. On the dichotomy in auditory perception between temporal envelope and fine structure cues (L). J Acoust Soc Am. 116:1351–1354.

Zeng F-G, Nie K, Stickney GS, Kong Y-Y, Vongphoe M, Bhargave A, Wei C, Cao K. 2005. Speech recognition with amplitude and frequency modulations. Proc Natl Acad Sci U S A. 102:2293–2298.

Zhu S, Wong LLN, Chen F. 2014. Development and validation of a new Mandarin tone identification test. Int J Pediatr Otorhinolaryngol. 78:2174–2182.

Zion Golumbic EM, Ding N, Bickel S, Lakatos P, Schevon CA, McKhann GM, Goodman RR, Emerson RG, Mehta AD, Simon JZ, Poeppel D, Schroeder CE. 2013. Mechanisms Underlying Selective Neuronal Tracking of Attended Speech at a “Cocktail Party.” Neuron. 77:980–991.

Zoefel B, VanRullen R. 2015. Selective Perceptual Phase Entrainment to Speech Rhythm in the Absence of Spectral Energy Fluctuations. J Neurosci. 35:1954–1964.

Zoefel B, VanRullen R. 2016. EEG oscillations entrain their phase to high-level features of speech sound. NeuroImage. 124:16–23.

